# Numerical discrimination in *Drosophila melanogaster*

**DOI:** 10.1101/2022.02.26.482107

**Authors:** Mercedes Bengochea, Jacobo D. Sitt, Thomas Preat, Veronique Izard, Laurent Cohen, Bassem A. Hassan

**Author notes:** Corresponding authors: L.C. and B.A.H.

## Abstract

Sensitivity to numbers is a crucial and evolutionarily conserved cognitive ability. The lack of experimental models amenable to systematic genetic and neural manipulation has precluded discovering circuits required for numerical cognition. Here, we demonstrate that in a two-choice task *Drosophila* fruit flies spontaneously prefer sets containing more objects. This preference is determined by the ratio between the two numerical quantities tested, a characteristic signature of numerical cognition across species. Individual flies maintained their numerical choice over consecutive days. Using a numerical visual conditioning paradigm, we found that flies are capable of associating sucrose with numerical quantities and can be trained to reverse their spontaneous preference for large quantities. Finally, we show that silencing LC11 neurons reduces the preference for more objects, thus identifying a neuronal substrate for numerical cognition in invertebrates. This discovery paves the way for the systematic analysis of the behavioral and neural mechanisms underlying sensitivity to numerosity.

## INTRODUCTION

The cognitive capacity to make decisions based on numerical information is not restricted to humans. By estimating numbers, animals can dilute predation risks, increase predation efficiency^1^, and maximize food intake^2^. They can also perceive the number of social companions^3^ and better communicate with conspecifics^4^. Accordingly, sensitivity to numbers has been documented in a wide variety of vertebrate species from primates and other mammals^5,6^ to birds^7,8^, amphibians^9,10^ and fish^11,12^ (for reviews see^13–15^).

The combination of numerical tasks with simultaneous neural activity recordings allowed the exploration of the neural basis of brain functions for numbers (for review see^16^). For example, it has been shown that single neurons in the prefrontal and posterior parietal neocortices for humans^17,18^ and primates^19^ spontaneously respond to specific numerosity. Similarly, ‘number neurons’ have been reported in the telencephalic nidopallium caudolaterale of corvid^20^. In fish, there is evidence for broad activation of the caudal telencephalon during numerosity changes^21,22^. The complete understanding of how the brain computes numerical information will require not only the recording of brain areas but also the manipulation of specific brain regions during the execution of the numerical task. The study of the neurophysiological basis of numerical cognition in vertebrates has thus far proven to be experimentally difficult due to a combination of challenges of accessibility, complexity and lack of tools.

Insects have long been used to explore the neuronal computations of complex behaviors (for a review see^23^). Recent evidence shows that numerical skills provide an advantage in terms of fitness also to invertebrates^24–26^, suggesting that some form of numerical ability may have evolved in common ancestors of insects and mammals over 500 million years ago. Number judgments are thought to be important to increase reproductive opportunities in beetles^27,28^; to improve predation strategies in spiders^29,30^ and ants^31^; to estimate the distance traveled through step counting in desert ants^32,33^ and to enhance foraging strategies in bees^34^. Several tasks have been developed to document the numerical abilities of insects. The most common approach consists in simultaneously presenting two stimuli that differ in numerosity. Observations suggest that animals often prefer larger sets of items. For example, individual carpenter ants spontaneously discriminate between two piles of dummy cocoons^35^. Similarly, when simultaneously presented with different sets of simulated shelters, crickets spontaneously choose the set with the larger number of items^36^. Furthermore, it has been shown that honeybees have a spontaneous preference for multiple ‘flowers’ only in comparisons where the number 1 was the lower quantity and where the ratio between the lower and higher quantity was at least 1:3^37^. The numerical abilities of bees have also been demonstrated using associative learning paradigms upon extensive training (30 to 100 trials^38–43^).

Despite significant behavioral evidence of numerical tasks in invertebrates, the neuronal bases of this process remain unknown. Surprisingly, to our knowledge, numerosity perception has not been studied in fruit flies (*Drosophila melanogaster*). The fruit fly model would offer an excellent experimental platform to uncover the genetic and neurobiological processes for numerical cognition mainly thanks to the availability of tools to label, manipulate and record activity of neurons and decipher their connectome. Recent studies suggest the existence of some form of magnitude judgment in *Drosophila*. For example, flies adjust their defensive behavior depending on the number of conspecifics in the group. In response to inescapable threats, flies’ freezing behavior decreases with increasing group size^44^. Another recent study showed that fruit flies also tune their social interactions to group size and density^45^. Whether such changes in behavioral strategies actually reflect a response to number or other continuous dimension of the stimuli is unknown.

Here, we report the first evidence of numerical discrimination in *Drosophila*. We show that flies display robust preference for more numerous sets of visual objects, and this discrimination between numerosities depends mainly on their ratio. Each fly deploys one of three dynamical behavioral patterns during numerical decisions, and sustains this behavioral strategy over time. Furthermore, spontaneous preference for large numbers can be modified by a single training trial of classical Pavlovian conditioning. Finally, we report that silencing LC11 neurons causes a decrease in numerical preference. These findings thus identify the first componant of the neuronal circuitry required for robust numerical discrimination in *Drosophila*.

## RESULTS

### *Drosophila melanogaster* show a spontaneous preference for larger numbers of items

To evaluate whether flies show spontaneous preference between sets of objects that differ in numerical size, we modified the classical Buridan paradigm^46^. Individual flies were placed in a circular arena and presented with 2 unreachable fixed high contrast visual targets situated at opposite locations (**Figure 1A**). Flies walked back and forth between the two targets while a camera recorded their movements from above for 15 minutes, allowing for the extraction of behavioral parameters. The two targets consisted of different numbers of stripes, squares or discs. We measured the amount of time each fly spent near a ‘preference zone’ (i.e. occupancy) in the vicinity of each stimulus (**Figure 1B**), and used the difference of durations divided by their sum as a preference index (PI). The PI thus ranged from -1 to +1, with a PI of 0 indicating the absence of preference.

**Figure 1:**
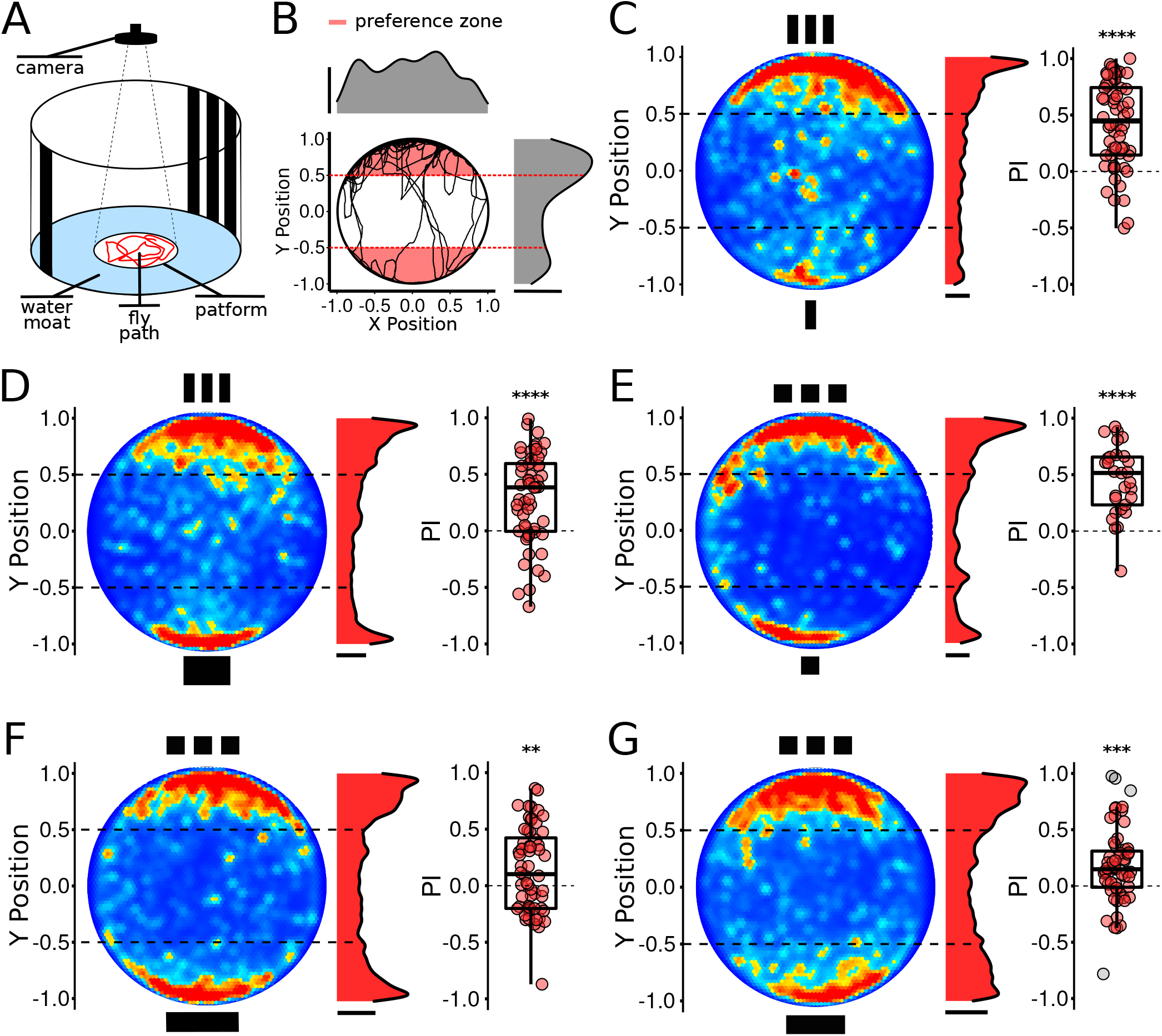
*Drosophila melanogaster* spontaneously prefer larger amounts of items in a 1vs3 choice test. **A**. Schematic of the experimental setup. We tested the spontaneous preference of individual flies between two unreachable visual number sets. A camera recorded the fly’s walking path (red line) from above for 15 minutes. **B**. Representative example of a fly’s individual walking path. Red dashed lines illustrate y-limits of the preference zone (red area) for each stimulus. Kernel density plots on the top and right of the platform denote the position permanence of the fly along x and y axes respectively. **C-D**. Canton S flies at population level show a significant spontaneous preference for three stripes over one stripe. Left: Heatmap illustrates the relative frequency of the fly location at each position of the platform (red denotes high frequency permanence while blue denotes low frequency). Middle: Kernel population y-density plot. Right: Mean population preference index is significantly different from chance preference (n=60, PI= 0.42±0.38, p=5.9e-09, Wilcoxon signed rank test). Control for overall area occupied (n=56, PI= 0.29±0.40, t_(55)_=5.41, p=1.4e-06, One sample t-test). **E-G:** Flies kept their preference for more objects when tested with arrays of squares. Flies’ performance in a 1vs3 squares contrast (n=29, PI= 0.45±0.31, t_(28)_=7.92, p=1.27e-08, One sample t-test). **F**. Control for overall area occupied (n=60, PI= 0.14±0.38, p=8.7e-03, Wilcoxon signed rank test). **G**. Control for total dark area (n=59, PI= 0.18±0.34, t_(58)_=4.00, p=1.8e-04, One sample t-test). Scale bar: 0.5 probability density. Asterisk indicates significance: **, p<0.01; ***, p<0.001; ****, p<0.0001.

We first evaluated the spontaneous PI for a set of 3 stripes versus a single stripe (**Figure 1A**). Flies stayed significantly longer in the preference zone of the arena corresponding to the set of three stripes (**Figure 1C, Figure S1Ai**). The same preference prevailed when the position of the stimuli was rotated 180° relative to the external environment (**Figure S1B1-2**) and no differences were found between sexes (**Figure S1C1-3**). However, this preference among sets differing in numerosity may result from a variety of potential confounds, including density, overall area of the display, dark area, and size of the individual shapes. Importantly, when flies were given a choice between three stripes and a single wide stripe occupying the same horizontal extension, they still preferred the three stripes (**Figure 1D, Figure S1Aii**) suggesting that the preference for the larger set was not due to the horizontal extent of the stimuli; nor to the total dark area, as the wide stripe had a larger dark area than the set of three stripes.

To test these findings with different displays, we replaced stripes with squares. Flies confronted with two single squares (1vs1) showed equal preference (**Figure S1D**). However, similarly to our previous observations with stripes, flies confronted with 1vs3 squares preferred the more numerous set (**Figure 1E, Figure S1Aiii**). Again, control experiments demonstrated that the preference was not due to the horizontal extent of the set (**Figure 1F, Figure S1Aiv**), nor to the total dark area (**Figure 1G, Figure S1Av**). Moreover, the pattern of preference remained the same when using discs instead of squares (**Figure S1E**). Flies also showed a significant preference for the more numerous set in 1vs4 (**Figure S1F**) and 1vs2 (**Figure S1G**) squares tests.

These results may indicate that flies are either sensitive to numerosity or are able to distinguish a single item from sets of several items. To differentiate between these hypotheses, we tested flies in a 2vs4 squares contrast. We found that flies prefer the larger set (**Figure 2A, Figure S2A**). In this experiment stimuli were equated for density and square size, but not for overall area occupied and total dark area. We next used the same numerical contrast to run a set of control experiments manipulating different variables that co-vary with number (**Figure 2B**). First, to equate overall area and square size we increased the distance between the two squares such that the outermost edges of the two visual sets matched. Flies still showed a preference for the set of four squares (**Figure 2C, Figure S2B**). Next, we equated the total dark area (while also keeping the horizontal extension equal), by increasing the size of squares in the set of two. Consistent with our previous findings, flies tested in this condition showed a significant preference for the four squares (**Figure 2D, Figure S2C**). Finally, we equated both dark area and overall area (dashed line in **Figure 2E**), while moving from a linear to a rectangular array of stimuli, providing a significant challenge to the numerical preference. Remarkably, flies still favored the numerically larger set (**Figure 2E, Figure S2D**). In addition, we observed similar responses in settings using 2vs3 squares contrast; flies still showed a reproducible preference for the larger set when controlling for density, overall area, dark area and spatial arrangement of the objects (**Figure S2E-H**). Finally, the preference for 3 over 2 objects was maintained when using other shapes, including stripes and discs (**Figure S2I-J)**.

**Figure 2:**
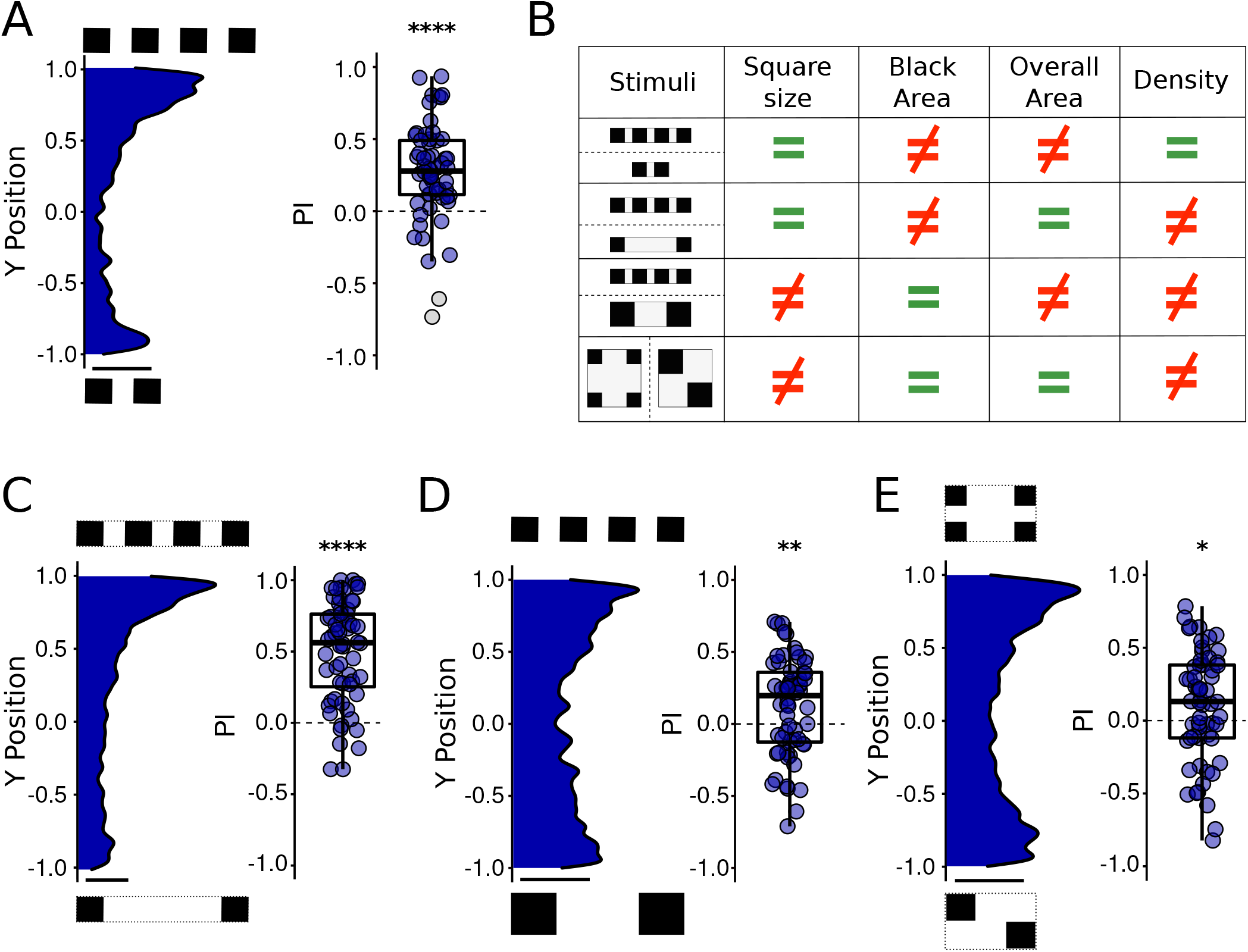
Flies spontaneously present a preference for more units in a 2vs4 numerical discrimination test, irrespective of non-numerical visual cues. **A**. Flies stayed longer near the 4-squares stimulus than near the 2-squares stimulus (n=60, PI= 0.27±0.33, t_(59)_=6.30, p=4.13e-08, One sample t-test). **B**. Table illustrating the various non-numerical features controlled in successive experiments. **C-E**. Series of experiments showing that the preference for numerically larger arrays is preserved when controlling for non-numerical cues. **C**. Control for overall area occupied (n=60, PI= 0.49±0.36 p=4.62e-10, Wilcoxon signed rank test). **D**. Control for total dark area and horizontal extension of the numerical sets (n=60, PI= 0.12±0.34, t_(59)_=2.74, p=8.1e-03, One sample t-test). **E**. Control for spatial distribution (n=60, PI= 0.10±0.38, t_(59)_=2.09, p=0.04, One sample t-test). Scale bar: 0.5 probability density. Asterisk indicates significance: *, p<0.05; **, p<0.01; ****, p<0.0001.

### *Drosophila* use the Approximate Number System to discriminate between numerosities

Animals rely on two different cognitive systems to process numbers in non-symbolic arrays, called the parallel individualization system or Object Tracking System (OTS) and the Approximate Number System (ANS). The OTS reflects the ability to simultaneously represent and track several items (usually up to 3 or 4)^47^, thus providing access to exact numerosity for only small arrays. In contrast, the ANS represents the approximate numerosity of sets of objects with no definite upper bound. Importantly, numerical discrimination by the ANS is governed by Weber’s law: The ability to distinguish between two stimulus magnitudes depends on their ratio. Hence, for example, it is easier to discriminate 4vs8 (a large, 1:2 ratio) than 4vs5; and moreover, it is easier to discriminate 5vs8 than 55vs58. Ratio-dependent discrimination has been observed in many species and is considered to be a characteristic signature of ANS^14,35,48,49^.

To test whether flies use the OTS or the ANS, we first asked whether they have a limit of numerical discrimination at 4 items, as observed in other insects^28,36,39,42,50^. We tested flies in a 3vs4 squares contrast, this time finding no preference (**Figure 3A, Figure S3A**). Importantly, this failure to discriminate 3 from 4 was not due to the similarity in total dark area of the two numerical sets: When the equivalent difference in total dark area was presented in a 2vs3 squares contrast, flies showed a significant preference for 3 squares (**Figure 3B, Figure S3B**), and this preference significantly differed from the response in the 3vs4 experiment (t_(108)_=-3.23, p=0.0016, Welch two sample t-test).

**Figure 3:**
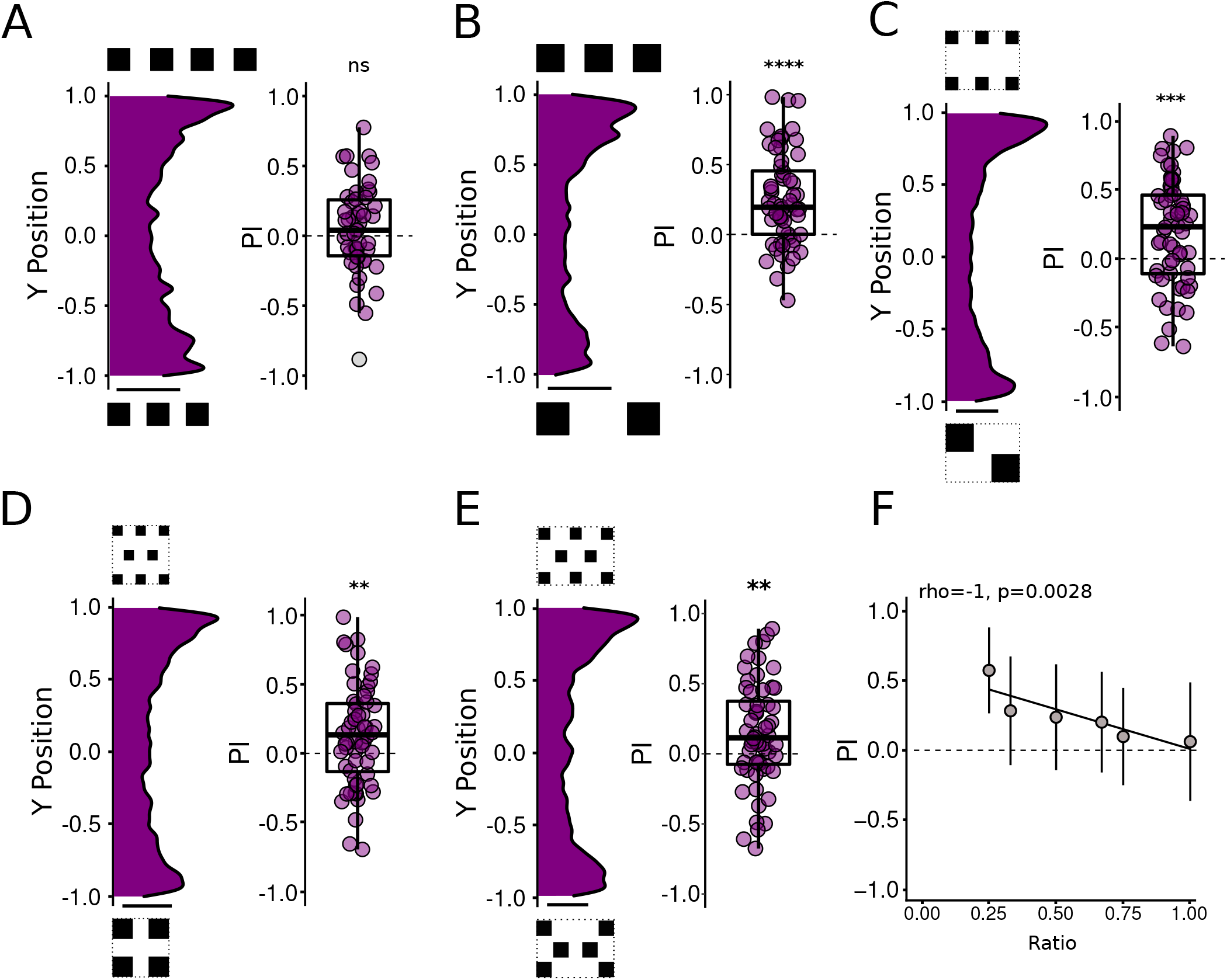
Flies use ANS to discriminate among numerosities. **A**. Flies tested in a 3vs4 squares contrast showed no numerical preference (n=50, PI= 0.05±0.32, t_(49)_=1.11, p=0.27, One sample t-test. **B**. Flies prefered 3 squares in a 3vs2 contrast with the same dark area as 3vs4 (n=60, PI= 0.25±0.33, t_(59)_=5.88, p=1.96e-07, One sample t-test. **C**. Flies preferred 6 squares in a 2vs6 contrast (n=60, PI= 0.16±0.40, t_(59)_=3.15, p=0.003, One sample t-test). **D**. Flies preferred 8 squares in a 4vs8 contrast (n=60, PI= 0.13±0.36, t_(59)_=2.82, p=0.007, One sample t-test). **E**. Flies preferred 8 squares in a 6vs8 contrast (n=60, PI= 0.14±0.37, t_(59)_=3.01, p=0.004, One sample t-test). **F**. Spearman correlation shows that discrimination accuracy decreases as the numerical ratio between quantities becomes closer to 1.0. Scale bar: 0.5 probability density. Asterisk indicates significance: *, p<0.05; **, p<0.01; ***, p<0.001; ****, p<0.0001; ns, not significant.

The failure to discriminate between 3 and 4 is compatible with both the OTS and the ANS systems. On one hand, flies may fail because numbers three and four are very close in terms of ratio (0.75), and cannot be distinguished by their ANS. On the other hand, flies may fail because the number 4 exceeds the capacity of their OTS. To distinguish between these two hypotheses we investigated whether flies can discriminate numbers larger than four. In these series of experiments, the set of squares were systematically equated for total dark area and overall area occupied. Flies consistently preferred larger numerosities in contrasts of 2vs6 (**Figure 3C, Figure S3C**), 4vs8 (**Figure 3D, Figure S3D**) and 6vs8 (**Figure 3E, Figure S3E**). These data suggest that flies may be using the ANS to perform numerical discrimination.

Knowing that flies were able to discriminate numbers higher than four, we wondered which parameter of the visual numerical stimuli best explains the numerical discrimination performance of the flies. Since it is impossible to control for all confounding variables within a single experiment, we implemented a global stepwise regression model (n=1361 flies, see **STAR Methods** for details) to examine the statistical relevance of each variable (total dark area, numerical ratio, total perimeter, larger numerosity and total overall area) resulting in a selection of variables that fit the numerical performance observed for flies. In terms of numerical parameters, the model predicts that flies primarily rely on the numerical ratio (t=-7.56, p=7.38e-14) but also on the larger numerosity (t=-2.59, p=0.01) to discriminate between sets of objects. To test the model’s prediction, we plotted the PI against the numerical ratio of all experiments and observed a significant negative correlation (R=-0.87, p=0.02, **Figure 3F, Figure S3F)**. This unbiased and global analysis confirmed that flies principally use numerical parameters to perform the discriminative task. Furthermore, the ratio-dependence of their performance strongly suggests that numerosity processing in *Drosophila melanogaster* is based on the ANS. Interestingly, the model also shows that the total dark area is a continuous variable that can influence preference (t=4.45, p=9.40e-06), reflecting the importance of having controlled for it in our experiments (**Figure 1G, Figure 2D-E, Figure S2G-H**).

### Flies display three individualized dynamical behavioral patterns for numerical discrimination

To understand the behavioral strategies used by flies to make ratio-based preference decisions, we performed cluster analysis of walking trajectories using all the spontaneous preference data, providing a typology of dynamic behavior during the preference task^51^ (**Figure 4**). We found three main clusters of behavioral patterns: 1. Flies that rapidly chose the larger set and remained there over most of the epoch (**Figure 4Ai**, green line in **Figure 4B**); 2. Flies that first hesitated between the two halves of the arena before eventually choosing the larger set (**Figure 4Aii**, orange line in **Figure 4B**); 3. Flies with no initial preference but later orienting weakly towards the smaller set (**Figure 4Aiii**, purple line in **Figure 4B**). Next, we identified the corresponding cluster for each individual behavior and studied the percentage of animals within each pattern. Animals tested with larger ratio contrasts (close to 0), predominantly display pattern 1, while animals tested with ratios closer to 1.0 displayed mainly patterns 2 and 3 (**Table S1**).

**Figure 4:**
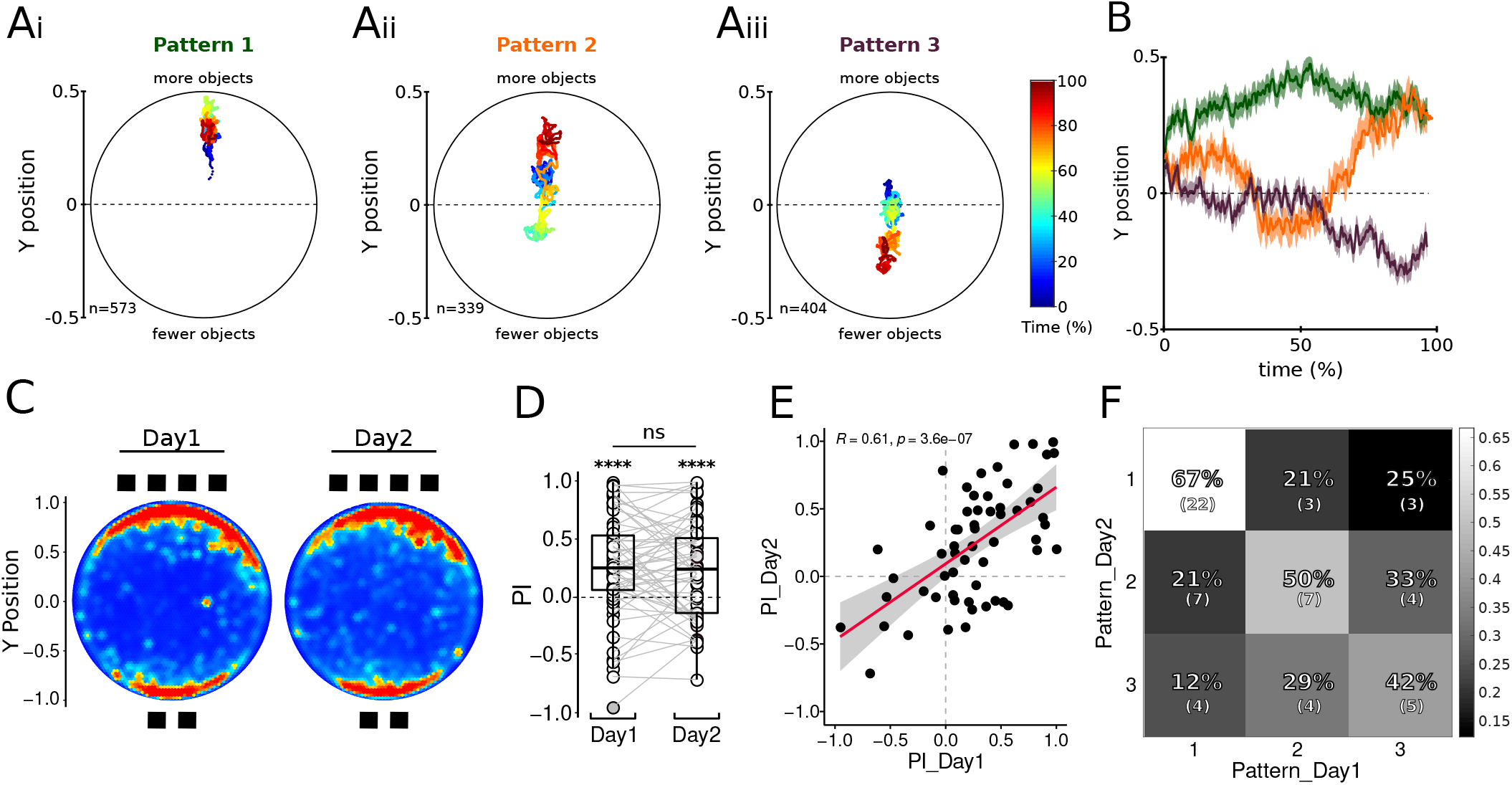
Flies display three main stable behavioral patterns of numerical preference. **A**. Flies display three main behavioral patterns. Plots show the mean trajectory for each pattern of behavior. Color code indicates the recording time (%). **i**. Pattern 1: Flies with stable preference for more items along all recording time; **ii**. Pattern 2: Hesitant flies that finally choose more items; **iii**. Pattern 3: Flies that show no preference at the beginning but then decide for less items. **B**. Y position over time for the three categories of behavioral preference. **C**. Population heatmap corresponding to each day. **D**. For each particular day flies prefered 4 squares against 2 (day1: n=59, PI= 0.26±0.44, t_(58)_=4.48, p=3.61e-05, One sample t-test; day2: n=59, PI= 0.24±0.42, t_(58)_=4.45, p=3.97e-05, One sample t-test). The performance is maintained between consecutive days (p=0.71; t_(58)_=0.37, Paired t-test). **E**. Pearson correlation plot showing a significant relationship between preference indexes of flies tested over two consecutive days. Gray shadow indicates confidence interval (95%). **F**. Matrix heatmap showing the relationship between patterns of behaviors in day1 and day2. The relative and absolute -between parentheses-number of animals with the specific relationship are indicated in each box.

To examine whether this reflected stable individualized tendencies, we repeatedly tested individual flies over two consecutive days using a 2vs4 squares contrast. As expected, on average flies chose the larger set on both sessions and the preference index did not differ between the two days (**Figure 4C-D**). Across flies, there was a strong positive correlation between the preference indexes in the two sessions (**Figure 4E**). Finally, we evaluated the stability of the temporal pattern using the clusters defined in the previous experiment. The fact that the clusters are defined on one dataset and tested on an independent dataset is an additional replication of the cluster validity. We assigned the trajectories of each fly on both days independently to the corresponding trajectory cluster (see **STAR METHODS**). We first validated that cluster 1 was predominant on both days (55.9% of flies on day 1, and 47.5% of flies on day 2). In addition we found that flies were highly consistent in their temporal pattern of preference in between days. In other words, the chosen pattern on day 1 was the most likely pattern on day 2 (**Figure 4F**, p=0.0014, Fisher’s exact test). These series of analysis suggest that flies have stable individual preferences when making number-based decisions.

### Spontaneous numerical preference can be modified by classical conditioning

Next, we further tested the cognitive numerical capacities of flies by asking whether they can associate a learned value to a specific numerosity and change their spontaneous preferences accordingly. Based on a novel appetitive classical conditioning paradigm, we endeavored to teach flies to reverse their spontaneous preference for larger numbers by associating sucrose (Unconditioned Stimulus, US) to the set containing smaller numbers. Individual wet-starved flies were placed in the arena and were trained in a single trial of 3 minutes. We paired a sucrose stimulus with the single square (Conditioned Stimulus, CS+) in a 1vs3 squares contrast (Trained group, TR). A control group (CT) was run in parallel by pairing the single square with water. For both groups, the set of 3 squares was water-paired (CS-). Two hours after training, flies from both groups were individually tested for five minutes in one non-reinforced test with the same stimuli. Flies that were trained to associate sucrose with the single square did not show a preference for the set of 3 squares in the testing session (**Figure 5A, Figure S4A**) in contrast to the control group (**Figure 5A, Figure S4B**). The preference differed significantly between the groups of flies trained with sucrose and the control group with no sucrose (**Figure 5A** right panel, **Figure S4A-B**), meaning that after the learning process trained flies spent on average more time near the single square.

**Figure 5:**
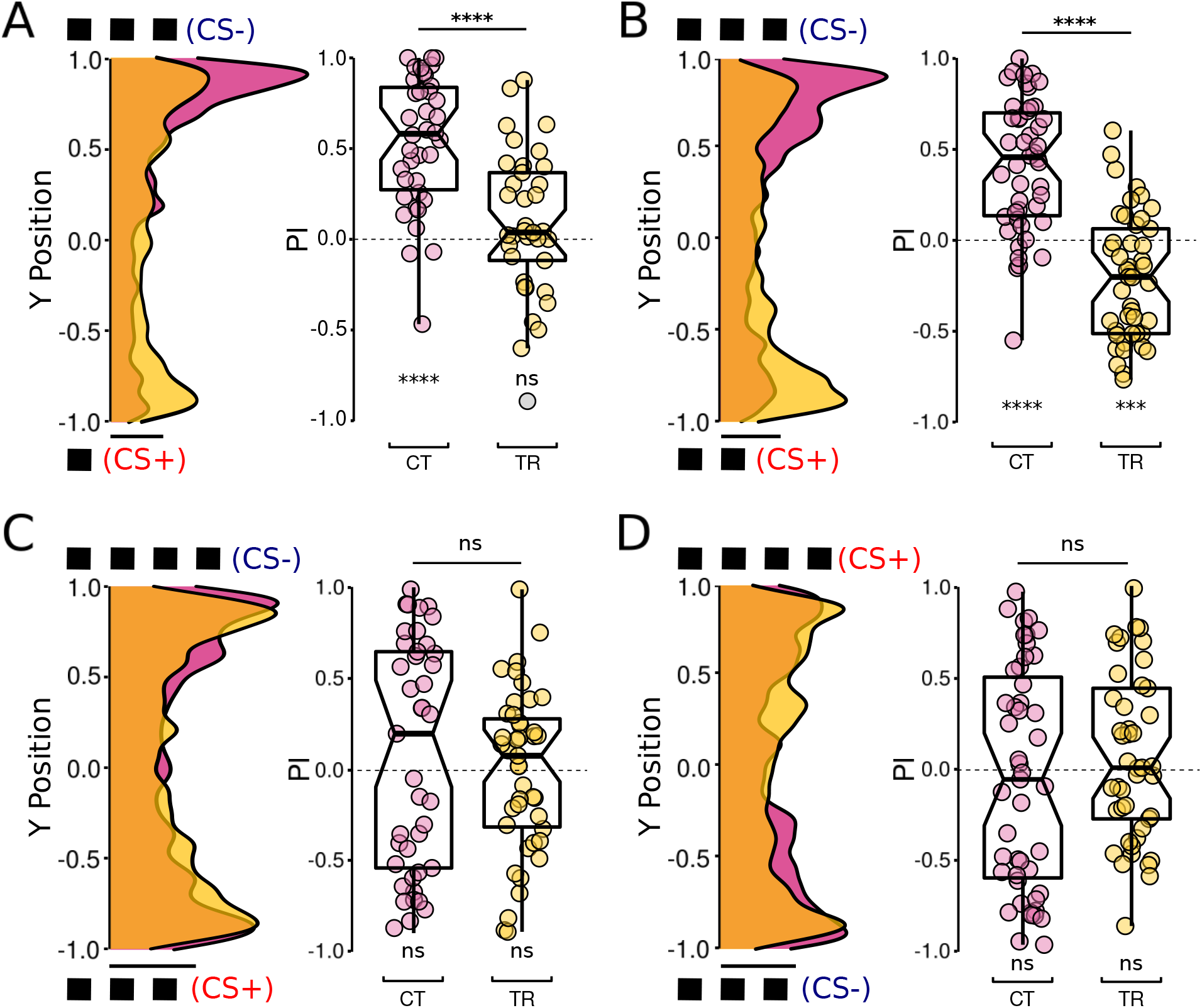
Flies’ numerical preference can be modified by associative conditioning. Wet-starved flies were trained to associate a non-preferred set of squares (i.e. smaller numerosities, CS+) to sugar (US), while the larger numerosity (CS-) was water-paired. The training consisted of one single trial three minutes long. Two hours later, the conditioned response was evaluated without the presence of sugar/water. **A**. Flies trained by pairing one square with sugar in a 1vs3 squares contrast showed no preference for three squares in the testing session. Left: Testing session kernel density plot for each group (Pink: Control group; Yellow: Trained group; Orange: overlap of the two curves). Control group spent more time in the 3 square zone of the platform while the trained group showed no numerical preference. Right: Unlike the control group (n=38, PI= 0.54±0.36, p=3.25e-09, Wilcoxon signed rank test), trained flies showed no preference for the set of three squares (n=37, PI= 0.08±0.36, t_(36)_=1.31, p=0.2, One sample t-test). Comparison between groups showed that flies trained to prefer one square significantly diminished the magnitude of their preference (p=1.8e-06, Wilcoxon rank sum test). **B**. Flies trained by pairing two squares set with sugar in a 2vs3 squares contrast showed inverse numerical preference during the testing session. Trained flies significantly preferred the smaller set of squares (n=45, PI= -0.20±0.34, t_(44)_=-3.93, p=2.9e-04, One sample t-test), opposite to the control group (n=42, PI= 0.39±0.36, t_(41)_=7.19, p=8.87e-09, One sample t-test; Comparison between groups: t_(84)_=7.9, p=7.94e-12, Welch two sample t-test). **C-D**. Flies trained with a numerical contrast they could not discriminate (i.e. 3vs4) did not show a change in their preference during the testing session. **C**. Flies trained to sugar-associate 3 squares did not show a zone of preference (n=41, PI= 0.008±0.44, t_(40)_=0.11, p=0.91, One sample t-test), same as the control group (n=40, PI= 0.08±0.64, p=0.33, Wilcoxon signed rank test; Comparison between groups: p=0.47, Wilcoxon rank sum test). **D**. Flies trained to associate 4 squares to sugar did not show a zone of preference in the testing session (n=45, PI= 0.08±0.47, t_(44)_=1.18, p=0.24, One sample t-test), same as the control group (n=46, PI= 0.03±0.60, p=0.71, Wilcoxon signed rank test; Comparison between groups: p=0.25, Wilcoxon rank sum test). Scale bar: 0.5 probability density. Asterisk indicates significance: ***, p<0.001; ****, p<0.0001; ns, not significant. TR: Trained group, CT: Control group.

The loss of preference observed after associative learning could be due to two different causes. First, it may be that the flies did not learn any association: during the testing session, they simply explored the arena in search for food. Second, it could be that flies did learn the association but that there was a competition between the learned response and the strong spontaneous preference for 3 squares compared to 1. To disentangle these possibilities, we trained the flies with a 2vs3 squares contrast that elicits a slightly weaker spontaneous preference, therefore decreasing the competition between the potentially learned response and the spontaneous response. Flies trained to 2 square-rewarded stimuli showed a switch in their preference during the testing session and preferred the smaller amount of items (**Figure 5B, Figure S4C**), in contrast to the control group (**Figure 5B, Figure S4D**). The preferences of the two groups were significantly different (**Figure 5B** right panel, **Figure S4C-D**). Moreover, the conditioned preference for the lower numerosity was significantly different when testing the flies in a 2vs3 or a 1vs3 contrast (p=0.0008, Wilcoxon rank sum test), suggesting a competition between the natural tendency to go for more items and the conditioned response.

Did flies learn to respond to numerosity, or to other variables of the visual displays? If the reversal of the preference is truly contingent on the numerosity of the trained stimuli, it should not occur in settings where flies are unable to distinguish stimuli based on their numerosity. We thus trained flies using a numerical contrast of 3vs4, which does not elicit a significant spontaneous preference. If flies had learned to respond to confounding variables - such as total dark area-we should find that they again develop a preference for the conditioned stimulus. In contrast, if flies learned to respond to the numerical variable, they should not learn when they cannot discriminate. We found that flies trained to prefer either 3 or 4 in a 3vs4 squares contrast show no preference during the testing session after training, similar to the control group (**Figure 5C, Figure S5A-D**). Finally, conditioned performance between animals trained to prefer 2 in the 2vs3 squares contrast and animals trained to prefer 3 or 4 were significantly different (2vs3-3vs4_”#$%&’_: p=0.013; 2vs3-3vs4_”#$%(‘_: p=0.003, Wilcoxon rank sum test). Together, these findings suggest that flies can learn to associate numerical sets with sucrose, and non-numerical aspects of the visual pattern (like overall area and total dark area) are not the properties of the stimuli that flies associate during the learning process.

### LC11 neurons mediate numerical discrimination

Knowing that fruit flies can discriminate among numerosities, we took advantage of the genetic toolkit that *Drosophila* offers to study which brain regions or neurons are necessary to this cognitive process. To our knowledge, there is no literature regarding the neuronal correlates of numerical abilities in insects. It has been proposed that this advanced cognitive skill could involve higher-order areas of the brain like the mushroom bodies or the central complex^49^. However, we found that silencing the mushroom bodies or the central complex using Tetanus Light Chain (TNT) did not affect the ability of flies to prefer the larger numerosity in a 2vs4 squares contrast task (**Figure S6A-D**).

Another set of studies proposed that the number sense is deeply ingrained into the primary sensory system^25,52^ as numerosity is mostly conceived as a primary sensory attribute^53^. Next, we tested the requirement of visual neurons involved in object orientation called medulla Dorsal Cluster Neurons (M-DCNs)^54^. However, we found that flies with silenced M-DCNs were able to discriminate between the visual sets, showing a preference for 4 squares (**Figure S6E-F**). In *Drosophila*, Lobula columnar neurons 11 (LC11) have been implicated in the adjustment of defensive behavior depending on group size, perhaps by making flies less sensitive to the movements of other flies^44^. We wondered whether LC11 is also involved in the detection of group size. Using the same 2vs4 contrast, we evaluated if LC11 neurons play a role in numerical discrimination by silencing their synaptic activity. We found that these flies were not able to discriminate between the stimuli (**Figure S6G-H**). LC11 neurons have also been shown to be required for small object response (∼10°)^55,56^, as were another set of LC neurons called LC10a^57,58^. However, in contrast to LC11, silencing LC10a neurons did not inhibit numerical discrimination (**Figure S6I-J**).

To further test the requirement of LC11 neurons in numerical discrimination and rule out the potential confounding effect of squares as small objects, we switched to large vertical stripes as it has been reported that LC11 neurons show weak or no responses to stripes and blocking them in fact enhances responses to elongated bars^55,56^. We found that silencing LC11 neurons (**Figure 6A**) reduced numerical discrimination in a 1vs3 stripes contrast (**Figure 6B, Figure S7A-B**) and abolished the preference in a 2vs3 stripes contrast (**Figure 6C, Figure S7C-D**). Importantly, LC10a silenced flies (**Figure 6D**) tested in 1vs3 stripes (**Figure 6E, Figure S7E-F**) and 2vs3 stripes contrasts (**Figure 6F, Figure S7G-H**) were able to discriminate under both conditions. In summary, these results show that silencing a specific type of visual neurons in the lobula of the fruit flies disrupts numerical discrimination.

**Figure 6:**
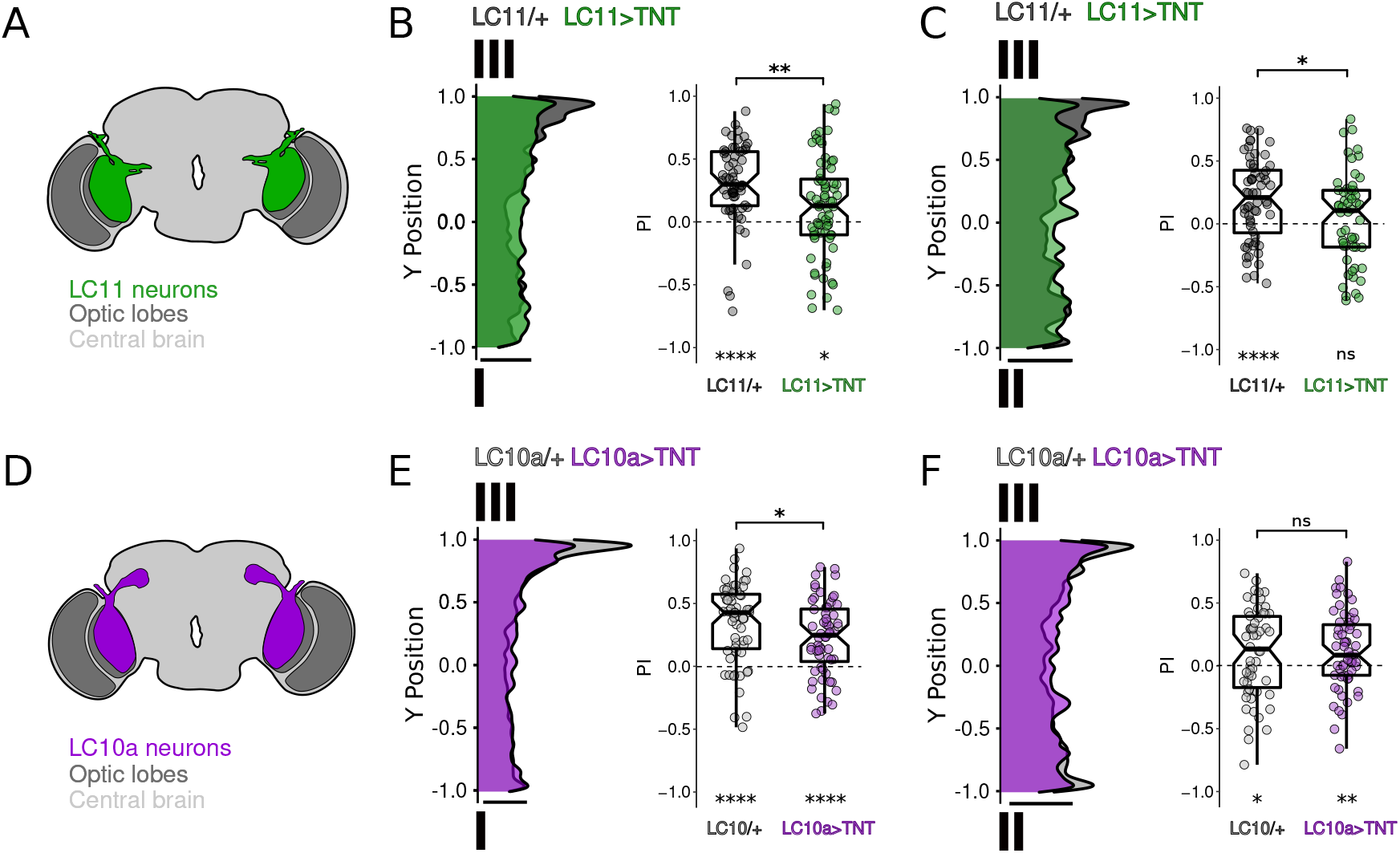
Silencing lobular columnar neurons 11 (LC11) diminish numerical discrimination. **A**. Schematic of the anatomy of LC11 neurons. **B**. Performance of flies with silenced LC11 tested in a 1vs3 stripes contrast. LC11/+: n=60, p=7.5e-08, PI= 0.30±0.32, Wilcoxon signed rank test; LC11>TNT: n=70, t_(69)_=2.55, p=0.013, PI= 0.12±0.39, One Sample t-test. Comparison between groups: p=0.0019, Wilcoxon rank sum test. **C**. LC11 silenced flies tested with a ratio closer to 1.0 (2vs3 stripes contrast) were unable to discriminate in opposition to control group. LC11/+: n=60, t_(59)_=4.43, p=4.1e-05, PI= 0.19±0.33, One Sample t-test; LC11>TNT: n=54, t_(53)_=0.81, p=0.42, PI= 0.04±0.36, One sample t-test. Comparison between groups: p= 0.032, Wilcoxon rank sum test. **D**. Schematic of the anatomy of LC10a neurons. **E**. Numerical discrimination performance of flies with silenced LC10a neurons in a 1vs3 stripes contrast (LC10a/+: n=60, p=1.19e-08, PI= 0.34±0.31, Wilcoxon signed rank test; LC10a>TNT: n=60, t_(59)_=5.96, p=1.52e-07, PI= 0.23±0.29, One Sample t-test; Comparison between groups: p=0.021, Wilcoxon rank sum test. **F**. Numerical discrimination performance of LC10a silenced flies in a 2vs3 stripes contrast (LC10a/+: n=58, t_(57)_=2.12, p=0.04, PI= 0.10±0.35, One sample t-test; LC10a>TNT: n=60, t_(59)_=2.88, p=0.005, PI= 0.11±0.30, comprison between groups, p=0.9, Wilcoxon rank sum test.) Scale bar: 0.5 probability density. Asterisk indicates significance: *, p<0.05; **, p<0.01, ****, p<0.0001; ns, not significant.

## DISCUSSION

### *Drosophila* as a model for numerical cognition

Numerical sensitivity is a crucial cognitive ability that is widespread across animal kingdom. Here, we developed the fruit fly *Drosophila melanogaster* as a new model for the neuroscience of numerical cognition. As a result, the powerful and versatile molecular, genetic and behavioral toolkit of *Drosophila* can now be exploited to unravel the neurobiological underpinnings of this highly conserved adaptive trait, while retaining individual animal resolution. Two main methodologies are described in the literature to study numerical cognition in different animal models: spontaneous discrimination tests and training procedures. In spontaneous choice tests, animals exposed to behaviorally relevant stimuli, such as high contrast objects in the case of fruit flies, reveal the preferences of animals for numerical sets. On the other hand, training protocols reveal an animal’s capacity to extract numerosities by assigning a positive value to a stimulus. We find that flies possess both capacities.

We demonstrate that flies spontaneously discriminate between sets of objects based on numerical quantity, showing a systematic preference for the more numerous sets with both small numerosities (1vs2, 1vs3, 1vs4, 2vs3, 2vs4) and larger numerosities (2vs6, 4vs8, 6vs8). This observation is reminiscent of a study showing that ants can discriminate between larger quantities of cocoons (2vs6, 2vs8)^35^. Further, we show that flies are able to discriminate numerical quantities independently of non-numerical continuous variables such as total dark area, total area, or object size. Moreover, their performance is ratio-dependent, a hallmark signature of the ANS, present in beetles^28^ and carpenter ants^35^. Interestingly, flies do not distinguish 3vs4 contrast, in line with findings in crickets^36^, spiders^59^ and honeybees^38^. However, our observations suggest that this may not reflect an upper limit of four, but rather a ratio limit.

A strict interpretation of Weber’s law, which is thought to govern the ANS, might suggest that when performing numerical discrimination flies, and likely other animals, predominantly use ratio. On the other hand, there is evidence that Weber’s law approximations may not always apply, and that humans and animals use various representations of numerical values that are combinatorially deployed depending on the characteristics of the task^60^. This is consistent with our observations that flies also consider continuous properties of the visual stimuli to discriminate among sets of visual objects. The outcome of the numerical performance observed in flies may be the result of the evaluation of numerical (ratio + larger numerosity) as well as non-numerical (total dark area) properties of the visual set presented. It has been shown that honeybees^56^ and crickets^36^ are able to perform relative numerosity judgements without knowing exactly the numerical value, rooting their estimation on non-numerical information. Animals likely use both types of parameters -as shown in fish^11^-depending on ecological needs. Future experiments like ‘equal-incongruent’ or ‘incongruent-opposite’ performed in bees^61^ will be needed to explore the importance of non-numerical variables in flies more deeply.

In addition to spontaneous discrimination, we find that flies are able to learn to associate a positive stimulus with numerical sets. Moreover, flies can reverse their preference for more items when fewer items are associated with sucrose after a single learning trail in a classical Pavlovian conditioning paradigm. This learning appears to be specifically associated with numerosity. Under conditions where flies cannot discriminate (3vs4), they fail to learn the position of the sucrose. This is despite the two sets differing in non-numerical variables such as total dark area and total area occupied, that could in principle be learned. Importantly, a similar discrepancy in dark area and total area occupied did not prevent flies from learning the association in a 2vs3 contrast, where the flies are able to discriminate numerosities. This supports the conclusion that the main criterion for learning under our single training session conditions is the numerical ratio.

Visual learning in adult flies has been extensively studied using operant conditioning in the flight simulator assay and in freely walking flies with single-fly and *en masse* approaches^62-63^. Here, we established a new behavioral paradigm for visual classical conditioning in freely walking individual adult *Drosophila*. Our protocol with a single training trial of three minutes generates a visual memory that lasts at least 2 hours allowing the study of short-term visual memories. It remains to be analyzed whether flies can form long-term numerical memories. In addition, it will be interesting to explore the range of flies’ numerical cognitive abilities -such as the aptitude to extract numerical rules-in flies using more extensive protocols^38–41^.

### Stability of numerical preference

One of the great advantages of fruit flies as a cognitive model is the possibility to study behavioral individuality in very large numbers of single flies. By analyzing the temporal dynamics of the numerical preference at individual fly resolution we show that flies display three main behavioral patterns of numerical preference. These categories allowed us to study the temporal stability of how flies make their numerical decisions. In a prior study^54^ we established a link between variability in the brain visual system of flies and the emergence of individuality of animal behavior. We find that individual flies show temporal consistency in their numerical choice. Future studies connecting neuronal morphology variation with the different numerical categories of behavior described here will allow us to unravel the neural bases of numerical preference. Moreover, it would be interesting to see whether the progeny of animals with specific numerical traits give rise to a behaviorally homogeneous population, or reproduce the population variance, as is the case with object orientation^54^.

### Number processing neural circuitry

A major quest in the field of numerical cognition is to identify neural pathways required for it. The availability of tools to selectively manipulate specific neuronal populations allowed us to test a small number of candidate neuronal subtypes involved in visual processing to provide proof of concept evidence for numerical processing in the fly brain. We describe the LC11 neuron as a neuronal type required for numerical discrimination, establishing the first starting point in the neuronal circuitry of numberical processing in invertebrates. When silencing LC11 neurons, flies showed a weaker numerical performance. Moreover, silencing a different type of lobula columnar neuron (LC10a) -which like LC11 also responds to small objects-or the central complex involved in visual navigation^64,65^ leaves the spontaneous numerical preference intact. This suggests not only some level of specificity for the role of LC11 in numerical processing, but also the emergence of numerical discrimination relatively early in the visual system. The advantage of using *Drosophila* as a model permits the use of a variety of sophisticated *in-vivo* techniques, including the combination of controlled behavioral tasks with simultaneous recordings of neuronal activity to investigate the brain circuitry associated with this ability. Future studies uncovering the neuronal circuitry of number-based judgment in fruit flies would be crucial to the understanding of number computation in insects and the role of LC11 cells in this regard. The comparative study of neuronal architectures across animals will unravel the evolutionary origin of number sense.

## ACKNOWLEDGMENTS

This work was supported by the Investissements d’Avenir program (ANR-10-IAIHU-06), Paris Brain Institute-ICM core funding, the Roger De Spoelberch Prize, an NIH Brain Initiative RO1 grant (1R01NS121874-01) (to B.A.H.) and The Big Brain Theory Program from the Paris Brain Institute (BBT.3400.COUNTINGFLIES, to L.C. and B.A.H). We thank all members of the Hassan lab for helpful discussions, Francois-Xavier Lejeune for statistical support and Dr. Carolina Rezával for comments on the manuscript.

## AUTHOR’S CONTRIBUTION

M.B, L.C. and B.A.H. conceived the study, designed the experiments, and wrote the manuscript. M.B. conducted all behavioral experiments and data analysis. J.D.S. performed cluster and statistical analysis. V.I. and T.P. provided expertise and helped write the manuscript.

## DECLARATION OF INTEREST

The authors declare no conflict of interest.

## EXPERIMENTAL MODEL AND SUBJECT DETAILS

*Drosophila melanogaster* were reared on a standard cornmeal/agar diet (8g Agar, 60g cornmeal, 50g yeast, 20g glucose, 50g molasses, 19ml ethanol, 1.9g Nipagin and 10ml propionic acid in 1L of water). Experimental animals were reared in groups up to 20 until 5 days old at 25°C in a 12/12 hour light/dark regime at 60% humidity. On day 5, the wings were cut under CO_2_ anesthesia. They were left to recover 48h within individual containers with access to fresh food before being transferred to the experimental set-up.

## METHOD DETAILS

### Behavioral arena

The behavioral arena used is a modification of the Buridan’s Paradigm^46^. The arena consists of a round platform of 119 mm in diameter, surrounded by a water-filled moat (**Figure 1A**). The arena was placed into a uniformly illuminated white cylinder. The setup was illuminated with four circular fluorescent tubes (Philips, L 40w, 640C circular cool white) powered by an Osram Quicktronic QT-M 1×26–42. The four fluorescent tubes were located outside of a cylindrical diffuser (Canson, Translucent paper 180gr/m2) positioned 145 mm from the arena center. The temperature on the platform during the experiment was maintained at 25°C.

### Visual stimuli

Accordingly with the particular experiment, different sets of black visual objects varying in width (*w*), height (*h*) and distance (*d*_*h*_ and *d*_*v*_ for horizontal and vertical distance respectively) were fixed inside the diffuser drum and opposite presented in the arena. The size of the dark area can be therefore described as: area = *w* x *h*. Apart from the stripes -that covered the whole vertical extension of the drum-the lower limit of the different objects were fixed to 30mm subtending an angular high of 11.7° from the middle of the platform. When the objects presented were discs, the diameter was fixed to 35mm. Generally, the distance between objects was 22mm subtending retinal sizes from 6.12° to 16.35° (8.6° in the center of the platform) except for the experiments where we controlled for the horizontal extension and the spatial distribution of the visual sets. **Table S2** describes the parameters of the different visual stimuli used in each experiment.

### Spontaneous numerical preference

To check for the spontaneous preference of the animals for stimuli of different numerical contrast we placed the flies on the arena for 15 minutes and tracked the walking path trajectory by using the software BuriTrack^46^. Each fly was tested once on a particular stimulus contrast. For half of the flies tested in each experiment, the drum was rotated 180 degrees to exclude any uncontrolled and systematic influence of other stimuli of the surroundings.

### Trajectory k-means clustering

To reveal the animals’ prototypical temporal patterns of spatial exploration across experimental conditions, we adapted a clustering approach based on k-means clustering, implemented through Matlab (Matlab R2016b and R2020a, The MathWorks, Inc.)^51^. Clustering was performed on the temporally resolved Y axis coordinate of each experiment/fly. Spatial trajectories across experiments were fisher transformed and temporally smoothened. All trajectory recordings were temporally aligned from the beginning of the experiment to a maximal length of 26500 samples (14.72 minutes). In this analysis when a fly’s trajectory could not be temporally aligned in more than 90% of the experiment (for example, if the fly made several jumps outside the arena) we discarded that given recording (45 flies rejected, 3.3% of the flies recorded).

Trajectories were iteratively partitioned into 2-16 clusters, in which each fly was assigned to the cluster with the nearest centroid trajectory. In detail, the procedure is as follows: (1) we define N (2 to 16) initial centroids corresponding to pseudo-randomly chosen individual trajectories (kmeans++ strategy, maximizing distance between initial centroids), (2) we compute the distance of each individual trajectory to the centroids by summing the euclidean distance -time point by time point-between each centroid and the given individual trajectory, (3) the individual trajectory is assigned to the cluster with the minimal distance, (4) the centroids are re-computed as the average -time point by time point-of all the trajectories assigned to that cluster, (5) the process is repeated until an stable assignment is determined for each trajectory, (6) the overall process is repeated 2500 times with different initial centroids and the iteration with minimal intracluster distance is selected.

The optimal number of clusters was determined using the elbow method. The intracluster distance showed a point of inflection for 3 clusters.

### Stability of numerical preference

To check whether the natural tendency of the flies was stable over time we tested the spontaneous preference of individual flies over two consecutive days. Flies were tested in a 2vs4 squares contrast for 15 minutes. After the first assay, each fly was placed back into their individual vial until the following day. Flies were tested at the same time of the day. Using the pre-defined clusters (learn on independent data, n=1316 flies), the behavior of each fly/trajectory was classed to the cluster with the nearest centroid trajectory. We did this analysis for each fly and day separately. Then, we calculated how many flies were classified in each cluster for each day and plotted the percentage of coincidence in the matrix.

### Associative learning experiments

For appetitive conditioning, flies were starved by placing them on individual vials with wet filter paper (EVIAN mineral water) 21hr before the training trial. The associative training consisted in one single trial of 3 minutes. Each individual fly was positioned in the arena and was trained once. As with the spontaneous preference test, the arena had two opposite sets of numerical cues. One set was used as sucrose-paired conditioned stimulus (CS+) and the other was used as unpaired conditioned stimulus (CS-). For each training trial, a round filter paper of 100 mm of circumference (Whatman CatNo 1001-110) was placed above the platform to support 2 to 3 drops of high concentrated glucose on the CS+ numerical set stimuli (1.5M, Sigma Aldrich G8270) and 2 to 3 drops of EVIAN water on the CS-numerical set condition. Between flies, one new fresh filter paper was used. In parallel, a control group was run in a second behavioral arena. Control flies had the same manipulation as trained ones but during the training trial we presented 2 to 3 drops of EVIAN mineral water on both sets of numerical stimuli. For both groups, flies that visited only one zone of the platform (CS+ or CS-) were discarded.

After the training trial, flies were placed back into their individual starved vial. The platform was cleaned with abundant distilled water and ethanol 70%. In order to test the short term conditioning, two hours later each fly was individually positioned into the arena for 5 minutes. During the testing session, one clean dry filter paper was positioned above the platform. As in the training, between flies the filter paper was renewed. Both groups were run in both set-ups and the position of the visual stimuli were rotated for half of the animals of each group.

### Data Analysis & Statistics

Statistical data was analyzed using R^66^. Transition plots were done as described before^46^. Briefly, the platform was divided in 60*60 hexagons and fly’s position raised the count of each hexagon by one in the arena. The scale starts at 0 (blue) and goes up till a value calculated by the 95%-quantile of the count-distribution (red). The arena was divided into three zones. To calculate the preference index (PI) we sum the density of passage of the hexagons within zones close to the visual stimuli (Red areas in **Figure 1B**) while the center part of the arena was not analyzed. Values indicate mean±SD. The preference index was calculated as

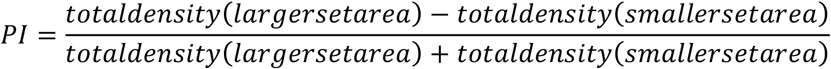

For the statistical analyses we first checked for normal data distribution using the Shapiro-Wilk normality test. Then, we chose the appropriate parametric or non-parametric test. For the spontaneous preference test we compared the PI against 0 (chance preference; 1 indicates preference for the larger set area and -1 indicates preference for the smaller set area) by using one-sample t-test or one sample Wilcoxon signed rank test. The null hypothesis was that choices would not be significantly different to chance expectation. We compared the mean of density occupancy for each area of the platform by using paired t-test or Wilcoxon Signed Rank Test. For the associative learning experiments, we statistically compared groups with the non-parametric Wilcoxon rank sum test. PI Boxplot: Each dot indicates the PI for each fly tested. Occupancy boxplots: Each blue dot corresponds to the permanence of a single fly in the area of the platform corresponding to the smaller set and each red dot corresponds to the permanence of the same fly in the larger set area. Boxplot elements: Center line: median; box limits: upper (75) and lower (25) quartiles; and whiskers, 1.5x inter quartile range; gray dots indicate outliers.

A forward stepwise linear regression was used to identify possible predictors of the outcome PI out of the following candidate variables: total dark area, numerical ratio, total perimeter, larger numerosity and total overall area. At each step, variables were added based on the p-value (<0.05) for an F-Test of the change in the sum of squared error that results from adding the term, and the p-value (>0.1) was used to remove a variable included in the final model.

---

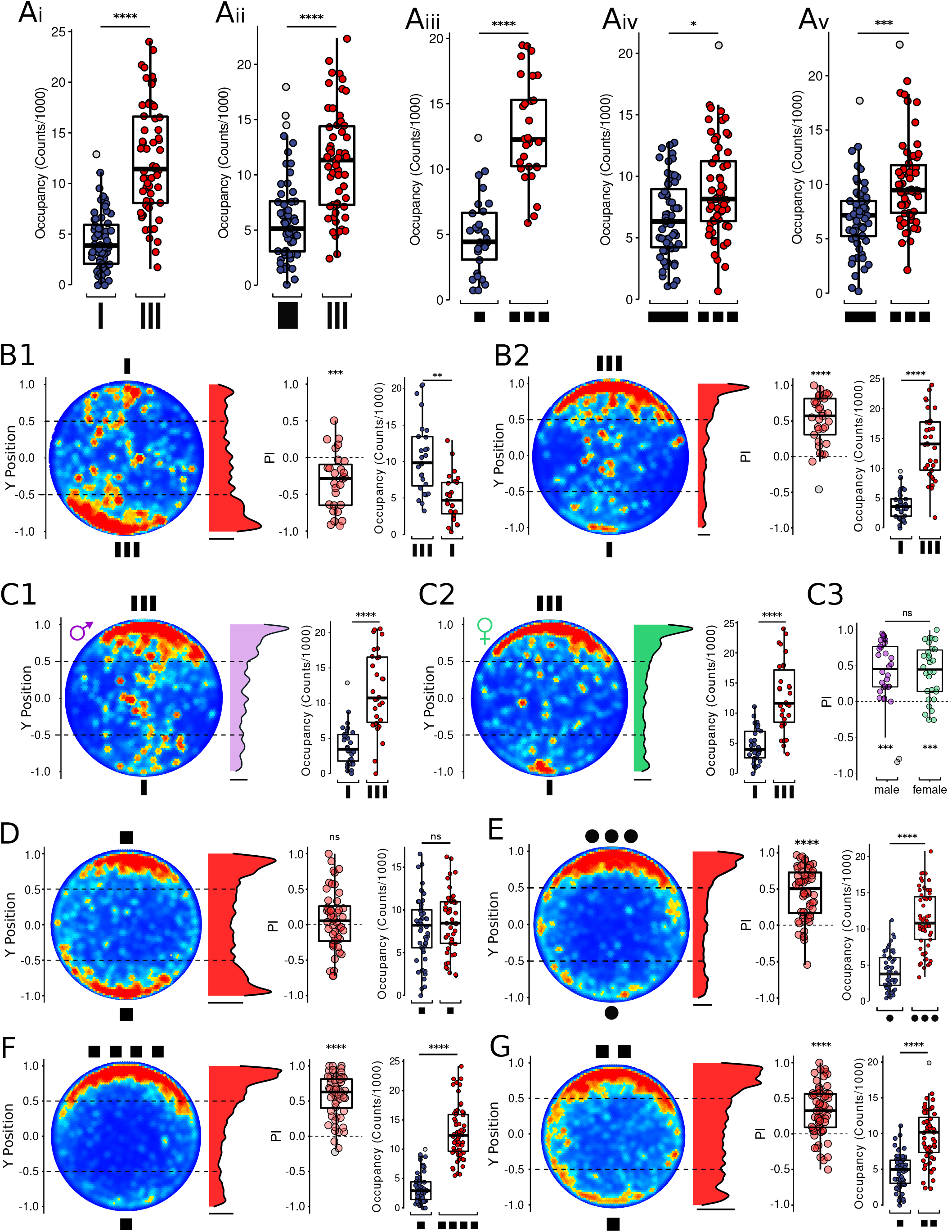

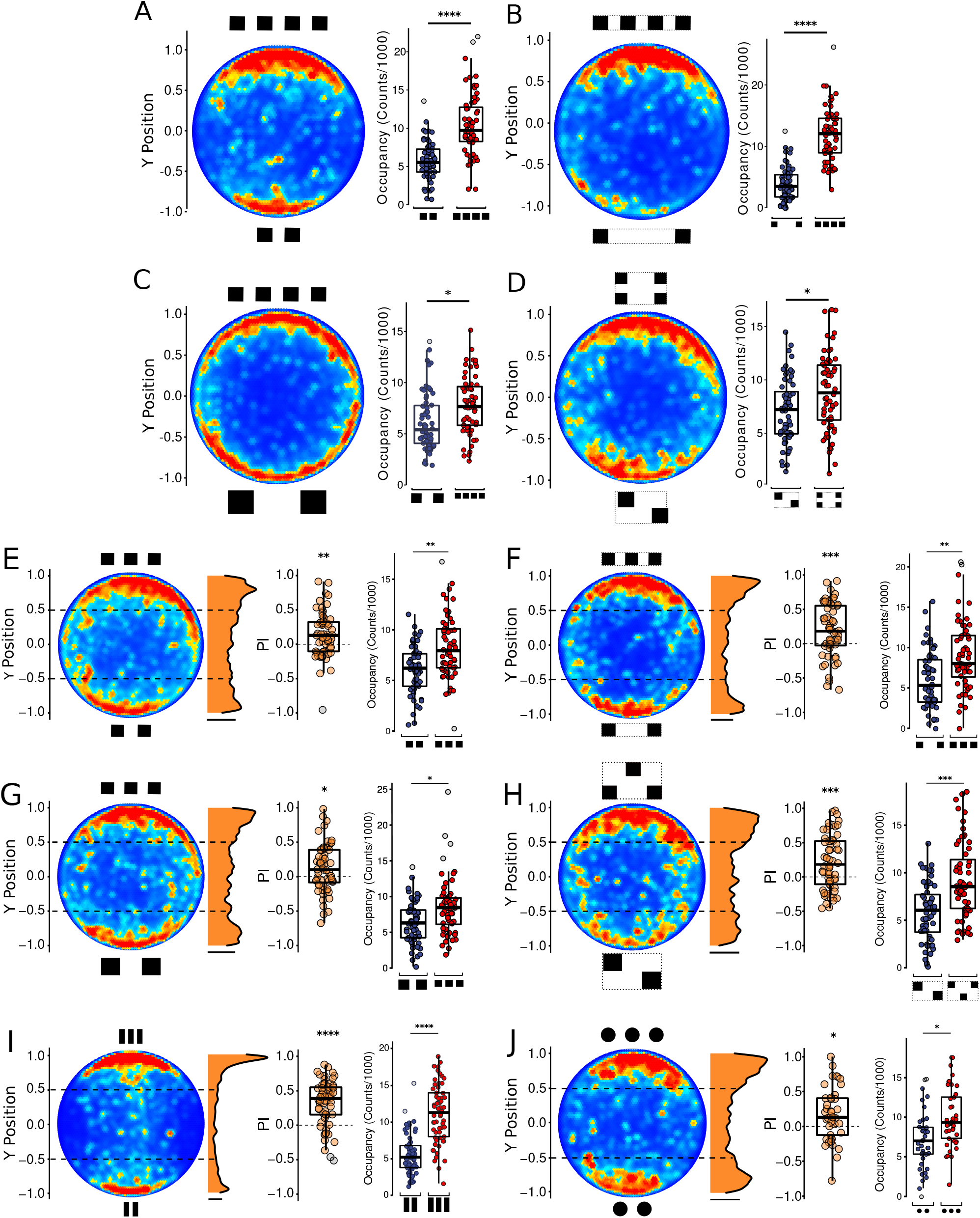

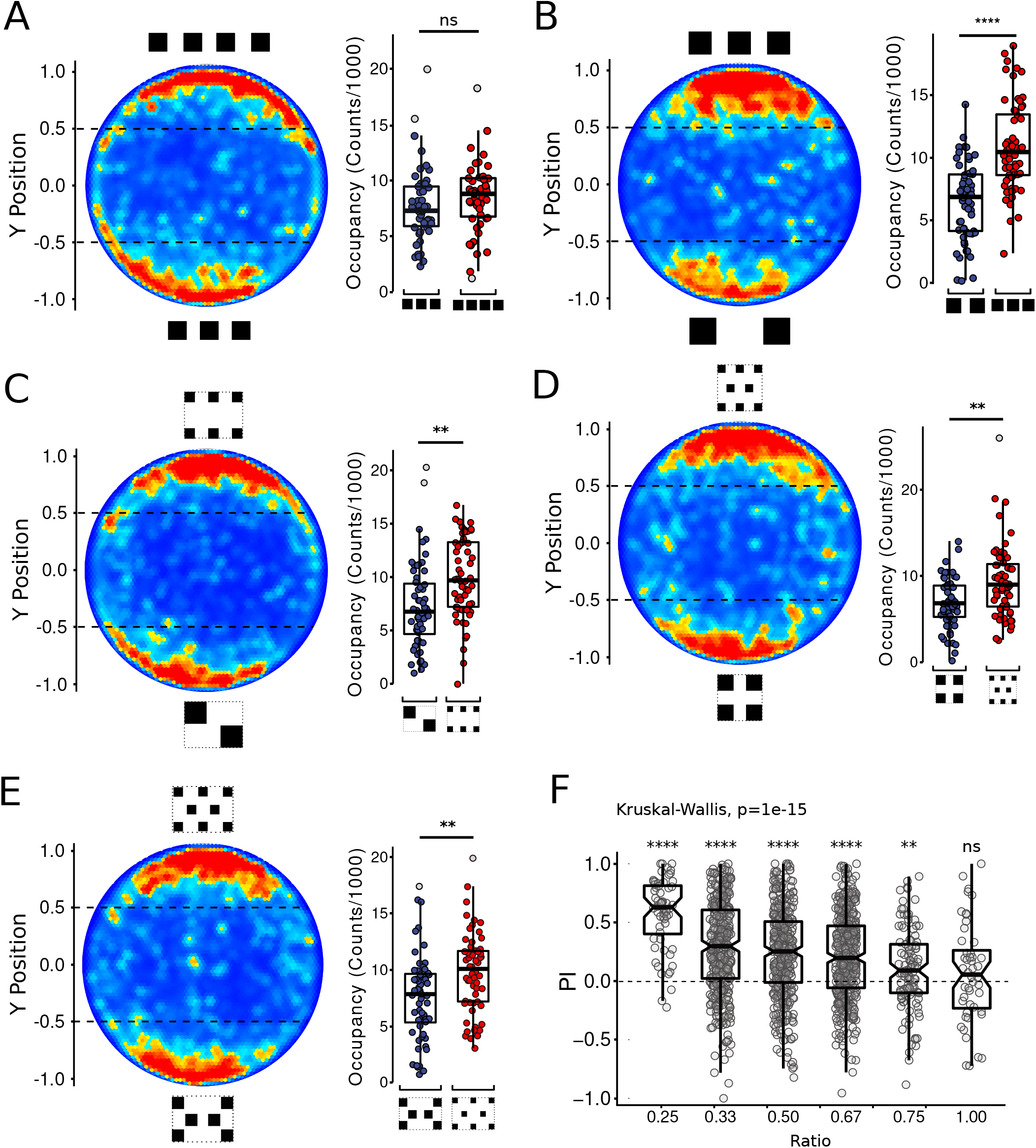

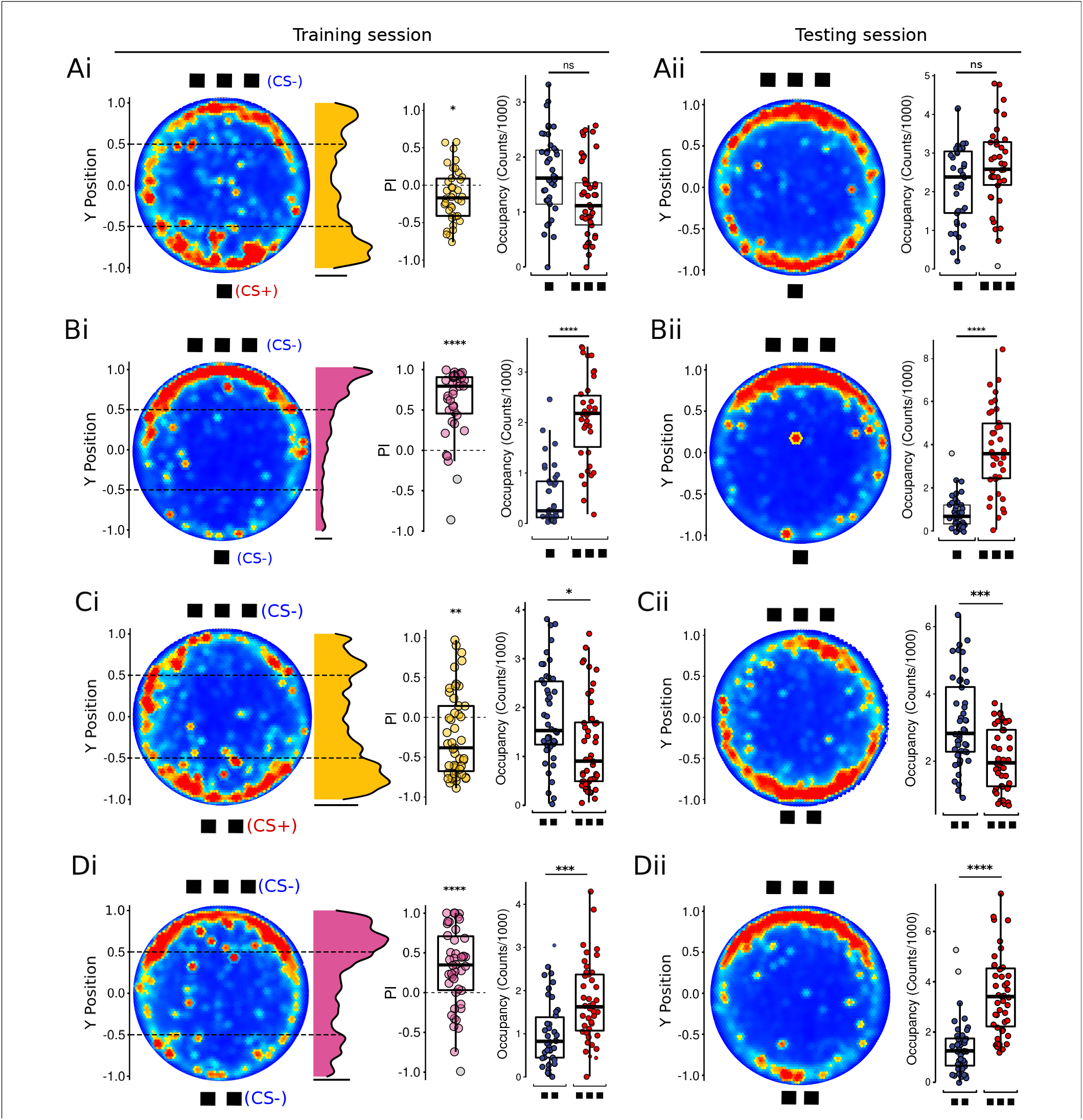

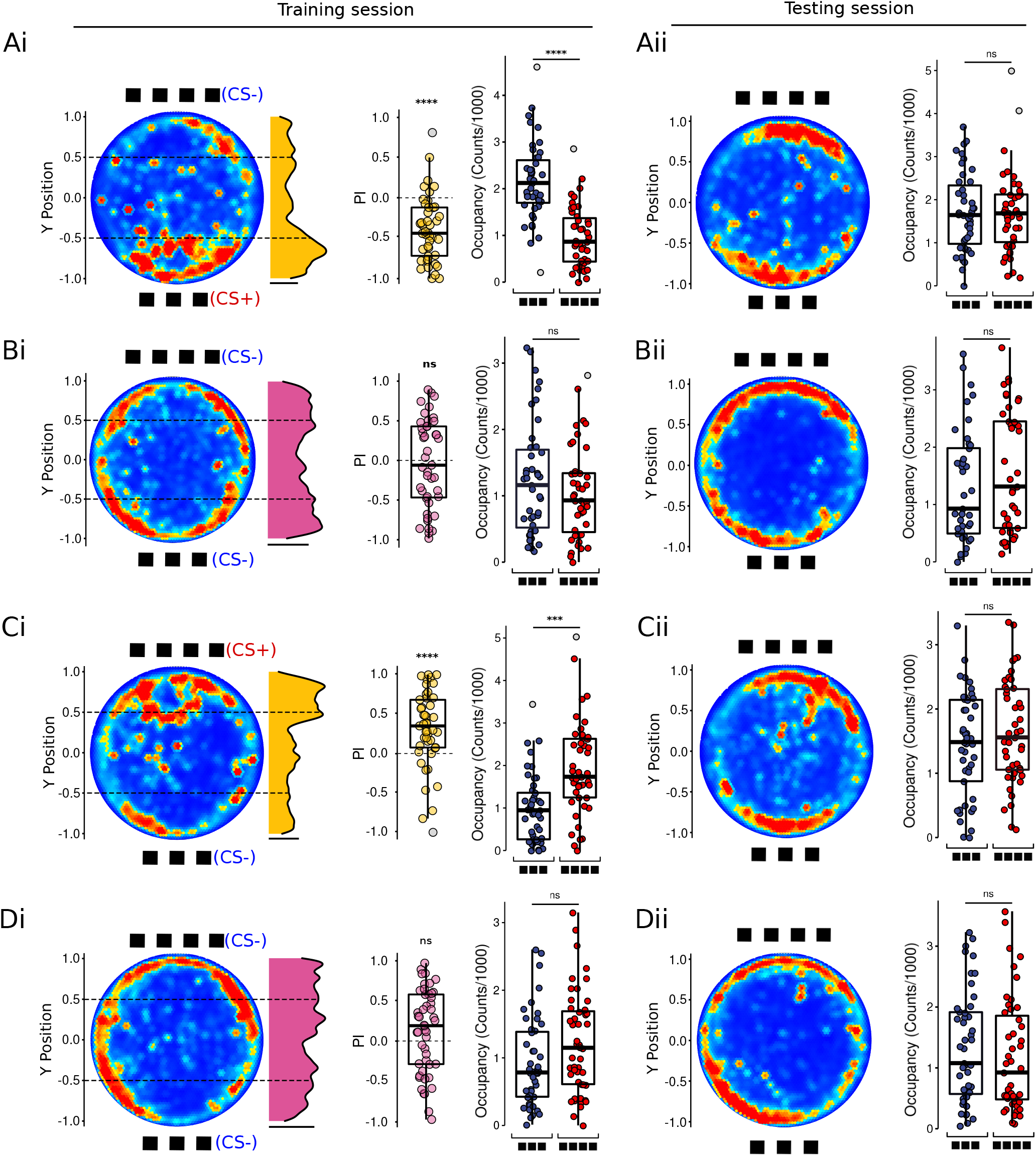

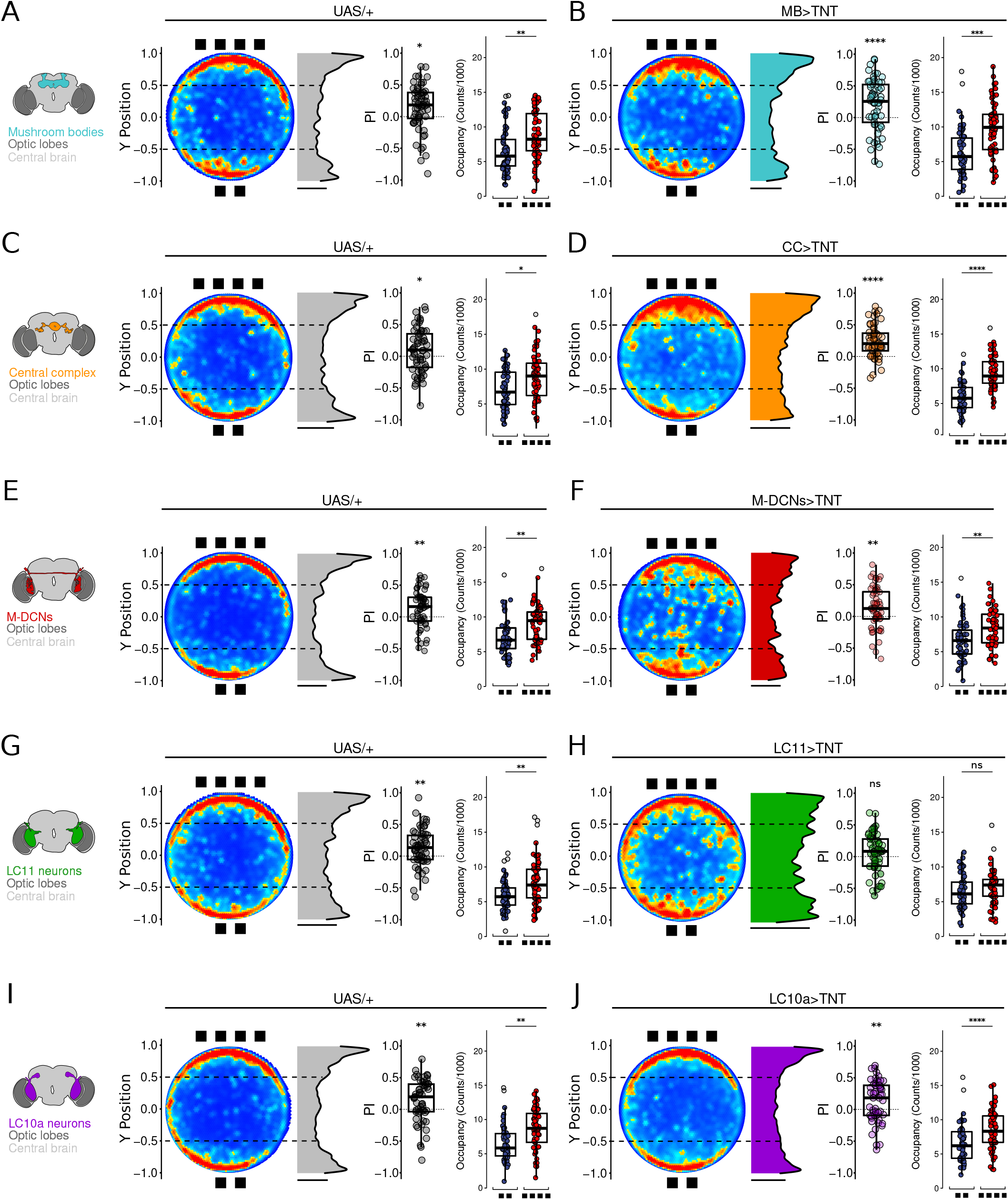

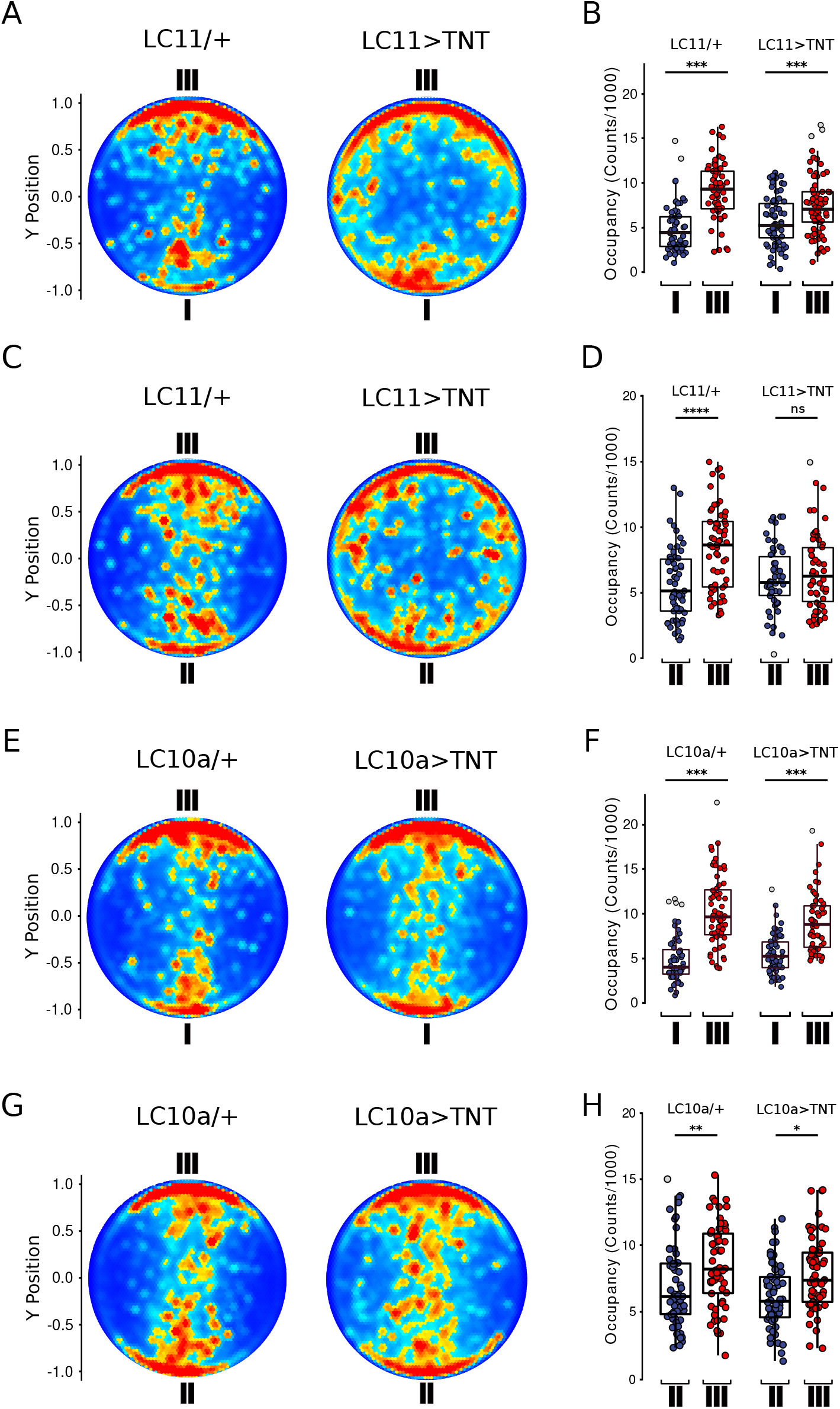

## Notes

### Competing Interest Statement

The authors have declared no competing interest.

